# Sensitized piRNA reporter identifies multiple RNA processing factors involved in piRNA-mediated gene silencing

**DOI:** 10.1101/2023.01.22.525052

**Authors:** Jordan Brown, Donglei Zhang, Wenjun Chen, Heng-Chi Lee

## Abstract

Metazoans guard their germlines against transposons and other foreign transcripts with PIWI-interacting RNAs (piRNAs). Due to the robust heritability of the silencing initiated by piRNAs in *C.elegans*, previous screens using *Caenorhabditis elegans* were strongly biased to uncover members of this pathway in the maintenance process but not in the initiation process. To identify novel piRNA pathway members, we have utilized a sensitized reporter strain which detects defects in initiation, amplification, or regulation of piRNA silencing. Using our reporter, we have identified Integrator complex subunits, nuclear pore components, protein import components, and pre-mRNA splicing factors as essential for piRNA-mediated gene silencing. We found the snRNA processing cellular machine termed the Integrator complex is required for both type I and type II piRNA production. Notably, we identified a role for nuclear pore and nucleolar components in promoting the perinuclear localization of anti-silencing CSR-1 Argonaute, as well as a role for Importin factor IMA-3 in nuclear localization of silencing Argonaute HRDE-1. Together, we have shown that piRNA silencing is dependent on evolutionarily ancient RNA processing machinery that has been co-opted to function in the piRNA mediated genome surveillance pathway.

## Introduction

To defend against invading nucleic acids, organisms must identify and silence foreign RNAs while preserving the expression of endogenous RNAs. When organisms fail to combat virus and transposon derived invasive nucleic acids, these elements can destabilize the genome and lead to widespread mutagenesis and infertility (1). One system animals have evolved to combat foreign nucleic acids is comprised of PIWI Argonaute and its associated small noncoding RNAs (piRNAs) (2–4). In *C.elegans*, piRNAs are able to initiate a robust silencing signal against mRNAs deemed as non-self (GFP, for example) which can be inherited for as many as 20 generations after the loss of the initiating PIWI protein complex (5). Because piRNA-initiated gene silencing is heritable independent of the initiation factors, piRNA reporters that rely on activation of a piRNA-silenced GFP transgene are biased to detect components that are required for maintaining and amplifying the already initiated silencing signal. For this reason, some of the requirements for the piRNA pathway in *C.elegans* likely remain unknown.

Here, we present a candidate RNAi screen in *C.elegans* using a reporter that relies on the previously characterized phenomenon that particular GFP transgenes have distinct propensities to be silenced when challenged with a perfectly complementary piRNA (6). We used this piRNA reporter to test for the involvement of snRNA processing factors, mRNA processing factors, germ granules components, and protein transport factors in piRNA-mediated silencing. We further characterized components from each of these categories to better understand how they mediate silencing in *C.elegans*. Our data suggest that the more recently evolved piRNA pathway utilizes highly evolutionarily conserved snRNA processing, pre-mRNA splicing, and protein import factors to carry out small RNA silencing, including the biogenesis of piRNAs and the regulation of Argonaute localization. Taken together, our results demonstrate that our piRNA reporter can identify various factors involved in distinct steps of the piRNA pathway. Our results also showed that proper piRNA-mediated transcriptome-wide surveillance relies on various regulatory factors that are not directly required for gene silencing.

## Results

### piRNA reporter detects loss of piRNA-dependent silencing factors

As described above, because piRNA targeting initiates a robust and stable silencing signal that persists even in the absence of the PIWI Argonaute protein PRG-1, previous reporter-based strategies have been biased to detect downstream components of the piRNA pathway (5,7). To remove this bias and increase the sensitivity of our screen, we aimed to establish a GFP reporter that is silenced in a piRNA dependent manner. We relied on previous observations that different GFP transgenes show distinct capacities to become silenced by the piRNA pathway (6). Our reporter strain contains two separate GFP transgenes with contrasting silencing propensities. One transgene, *cdk-1∷gfp*, becomes stably silenced when challenged by an artificial piRNA (integrated into endogenously encoded piRNA locus *21ur-5499*) that perfectly complements a 20nt site within the *gfp* coding sequence (8). A second transgene, *oma-1∷gfp*, cannot be silenced by a perfectly complementary piRNA (6). When we crossed these two lines to one another to obtain a strain that contains both GFP transgenes as well as the perfectly complementary piRNA, both transgenes are silenced. We hypothesized that because this strain contains one transgene that is vulnerable to piRNA silencing and another that resists piRNA silencing, then perhaps the silenced state of GFP is less stable and easily perturbed. To test this hypothesis, we introduced a frameshift mutation to the sole PIWI Argonaute *prg-1* using the CRSIPR/Cas9 system. We found that both transgenes became expressed at the first generation of *prg-1* homozygosity (F2 generation) after injection (Fig 1A). Therefore, this reporter can successfully detect loss of *prg-1*, suggesting that a screen using this sensitized reporter would be sensitive to loss of upstream factors in the piRNA pathway.

**Figure 1.**
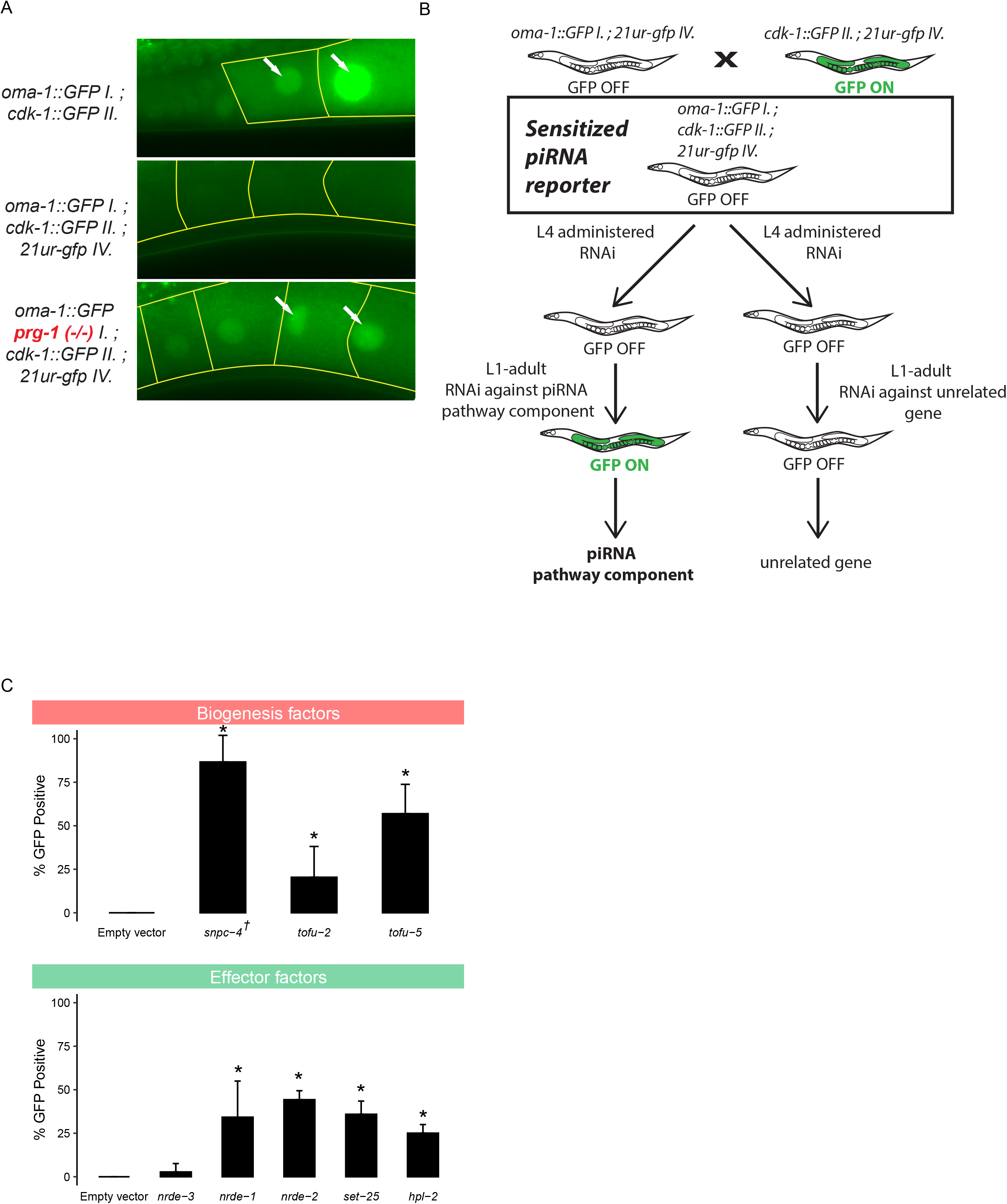
piRNA reporter detects loss of biogenesis and effector factors. A) Fluorescent micrographs show the expression of GFP transgenes in oocyte cytoplasm and nuclei in the absence of a triggering piRNA or *prg-1*. B) Schematic shows the screening strategy using the sensitized piRNA reporter. C) GFP expression in the piRNA reporter in indicated RNAi knockdowns. † indicates L1 to adult same generation RNAi treatment. Asterisks indicate significant reporter activation, see Methods for details.

Although many factors in the *C.elegans* piRNA pathway have been characterized, there are likely unknown components of this pathway that have not yet been discovered due to the use of less sensitive reporters in genetic screens. We therefore sought to use our sensitized reporter to identify components involved in the pathway both at the piRNA biogenesis level and at the effector level (Fig S1). We used a candidate approach by selecting specific factors to knockdown using RNAi (Fig 1B). To ensure that this approach can successfully identify the loss of factors at both of these levels, we performed RNAi against known piRNA biogenesis and effector components. We saw that depletion of piRNA biogenesis factors *snpc-4*, *tofu-2*, and *tofu-5* resulted in significant GFP expression in the reporter, as did effector factors *nrde-1*, *nrde-2*, *set-25*, and *hpl-2* (Fig 1C). As expected, we failed to observe significant GFP expression in the reporter when a somatic RNAi factor, *nrde-3*, was depleted with RNAi. These results suggested that our sensitized reporter can be used to detect loss of piRNA components using an RNAi based approach. The full list of candidates tested using our reporter can be found in Supplementary Table 1.

### RNAi screen identifies P granule factors, mRNA processing factors, and protein transport factors as piRNA silencing components

piRNA transcription in *C.elegans* is accomplished by RNA pol II. Since piRNA precursor transcription requires the SNPC-4 complex, which is also involved in initiation of snRNA transcription, we hypothesized that the Integrator complex, a multimeric snRNA processing complex required for snRNA 3’ end processing, may be involved in the termination of piRNA transcription. Our model is consistent with the recent report that termination of piRNA precursor transcription involves the Integrator complex (9,10). We performed RNAi against 9 different components of the Integrator complex in the sensitized reporter and found four of the components, *dic-1*, *ints-1*, *ints-9*, and *ints-11*, triggered GFP expression in the reporter following knockdown (Fig 2A). We failed to observe GFP expression following knockdown of *ints-4*, *ints-5*, *ints-7*, *ints-12*, or *ints-13* despite the gross morphological phenotypes resulting from most of these treatments. As individual knockdown of *ints-1*, *ints-2*, *ints-4*, *ints-5*, *dic-1*, *ints-7*, *ints-8*, *ints-9*, or *ints-11* has been shown to disrupt snRNA 3’ end processing in *C.elegans* (11), yet our results indicate that only individual knockdown of *ints-1*, *dic-1*, *ints-9*, or *ints-11* leads to defects in piRNA silencing, our finding suggests that a subset of the Integrator complex could be functioning at piRNA loci in a fashion distinct from its function at snRNA loci and coding genes. It has been shown recently that the mammalian Integrator complex does indeed form structurally distinct subcomplexes (12). Although a subcomplex composed of *ints-1*, *dic-1*, *ints-9*, and *ints-11* was not reported, our results suggest that these four components promote piRNA biogenesis.

**Figure 2.**
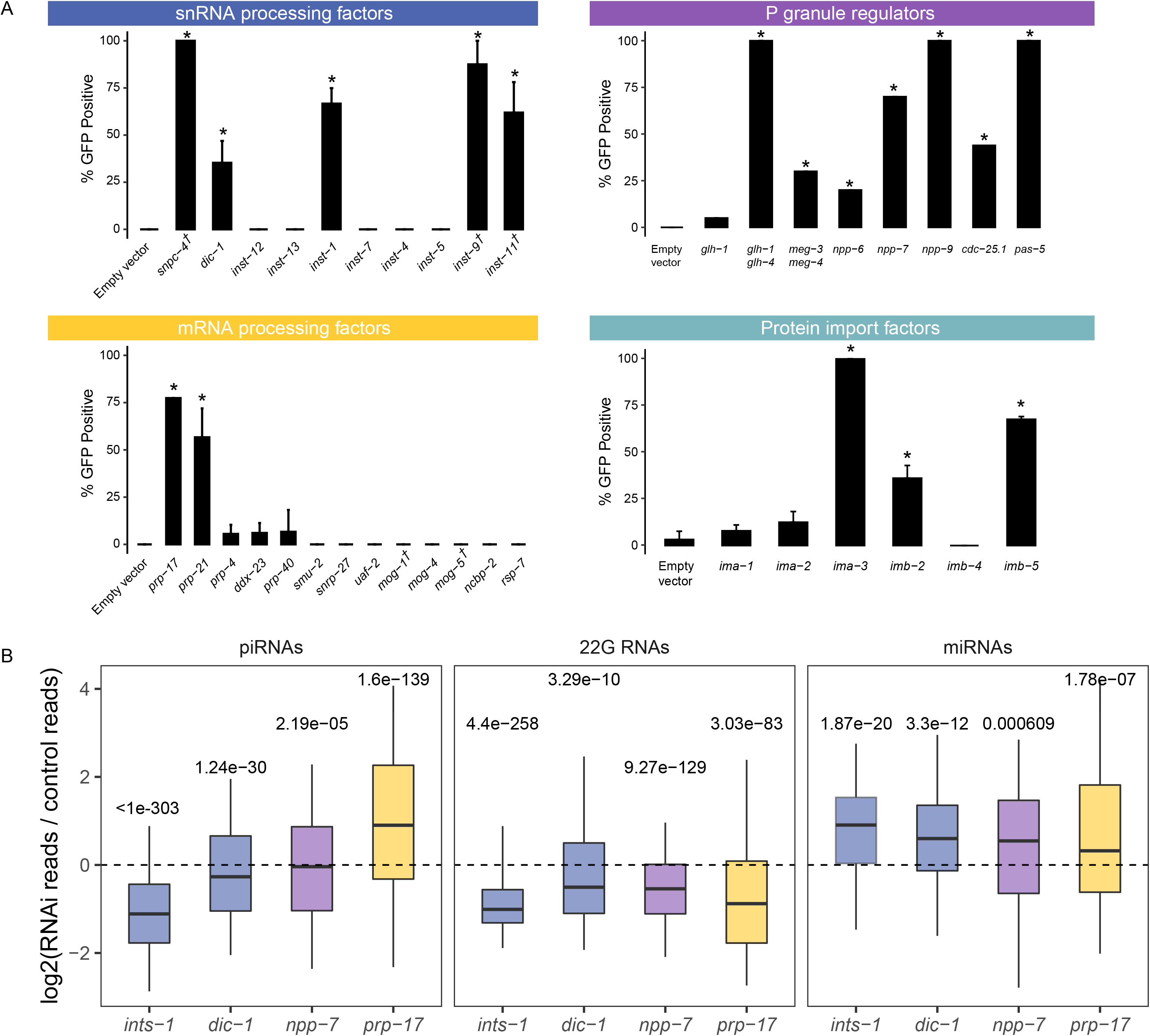
Loss of mRNA processing factors, snRNA processing factors, protein import factors, and P granules components trigger piRNA reporter activation. A) GFP expression in the piRNA reporter in indicated RNAi knockdowns or mutants (for *glh-1 glh-4* and *meg-3 meg-4*). † indicates L1 to adult same generation RNAi treatment. Asterisks indicate significant reporter activation, see Methods for details. B) Small RNA fold changes in the indicated RNAi knockdowns relative to empty vector control knockdown.

piRNA pathway components are enriched in phase-separated perinuclear organelles called P granules. Previous studies have shown that when P granules are lost, RNAi is compromised (13). It remains unclear, however, whether specific P granule regulators are important for gene silencing or whether the structures themselves are indispensable. We addressed this question by treated the sensitized reporter with RNAi against genes which have previously been shown to be enriched in or important for the formation of P granules (14). We found that loss of both *glh-1* and *glh-4* or *meg-3* and *meg-4* led to GFP expression in the piRNA reporter (15). In addition, knockdown of nuclear pore components *npp-6. npp-7*, and *npp-9*, phosphatase *cdc-25.1*, and proteasome component *pas-5* all led to GFP expression in the reporter (Fig 2A). The specific role of *npp-6/7* in promoting piRNA silencing is further explored below.

Pre-mRNA splicing has recently been shown to act as a signal for endogenous RNAi pathways in *C.elegans* (16–18). In addition, knockdown of some splicing factors leads to defects in P granule formation (14). We therefore wondered whether loss of particular splicing factors would trigger GFP expression in our sensitized reporter. We found that of the 13 splicing factors we tested, only loss of the second step factors *prp-17* and *prp-21* triggered GFP expression (Fig 2A), despite the gross morphological phenotypes resulting from treatment with most other RNAi constructs. This finding suggests some specific pre-mRNA splicing factors contribute to piRNA silencing, consistent with reports that splicing factor EMB-4 is essential to piRNA-mediated silencing for reporters containing multiple introns (17,18).

piRNA targeting triggers a silencing signal that relies on siRNAs to function both with secondary Argonaute proteins called WAGOs (Worm-specific ArGOnautes) in the cytoplasm and in the nucleus. The RNA-dependent RNA polymerases that manufacture these siRNAs, termed 22G RNAs due to their possession of a 5’ guanosine and 22nt length, all exist and function in the cytoplasm (19,20). Some of these 22G-RNAs associate with HRDE-1 (WAGO-9) Argonaute which is critical for the inheritance of gene silencing over generations (21). However, it remains unknown how HRDE-1 and piRNA-dependent 22G RNAs translocate into the nucleus in germ cells. We hypothesized that HRDE-1 binds 22G RNAs in the cytoplasm and subsequently translocates into the nucleus as has been shown with the nuclear PIWI Argonaute in *Drosophila* and the somatic nuclear Argonaute in *C.elegans* NRDE-3 (22,23). To identify the protein import machinery required for nuclear localization of HRDE-1, we depleted several Importin factors in our sensitized reporter and observed GFP expression following *ima-3*, *imb-2*, and *imb-5* knockdown, but not following *ima-1* or *ima-2* knockdown (Fig 2A). Since *ima-1* RNAi knockdown led to sterility but not piRNA reporter activation, this suggests that specific protein import machinery is essential for piRNA-mediated gene silencing (see more details on HRDE-1 nuclear localization below).

Because our sensitized reporter can detect piRNA pathway defects at any level of the pathway, such as piRNA biogenesis or production of 22G-RNAs (Fig 1C), we next sought to characterize the defects contributed by loss of the factors reported above. To accomplish this goal, we performed RNAi against a subset of these factors and sequenced the small RNAs from a population of depleted animals. While loss of *npp-7*, *prp-17*, *ints-1*, or *dic-1* all led to a reduction in 22G accumulation (Fig 2B), only RNAi of Integrator complex components *ints-1* and *dic-1* caused a reduction in piRNAs as well. Reduction of both piRNA and 22G-RNA levels is consistent with the Integrator complex’s involvement in piRNA biogenesis, as compromised piRNA production can also lead to reduced 22G-RNA accumulation as is the case in *prg-1* mutants (24). In contrast, the reduction in 22G-RNA but not piRNA levels suggests that *npp-7* and *prp-17* likely function as effectors in the piRNA pathway and do not function in piRNA biogenesis. We then characterized the defects in each of these cases further.

### Integrator complex resolves piRNA precursor 3’ ends and promotes the production of both type I and type II piRNAs

We found that loss of a subset of Integrator complex components caused GFP expression in our sensitized piRNA reporter, consistent with a recent report (9). We sequenced small RNAs from animals depleted for Integrator components *ints-1* or *dic-1* and observed a global reduction in mature piRNA production (Fig 2B and Fig S2A). In addition, piRNA precursor levels are strongly decreased following *ints-1* and *dic-1* knockdown (Fig S2B). In mammals and *Drosophila*, Integrator complex action at snRNA loci requires the SNAPc complex of proteins to bind the promoter of snRNA genes (25). The SNAPc homolog SNPC-4 in *C.elegans* is also required for piRNA biogenesis and interacts with piRNA promoters (26). We reasoned that if SNPC-4 recruits the Integrator complex to piRNA loci, then piRNAs that rely most on SNPC-4 for their accumulation should also rely on the Integrator. We found that this was the case for both INTS-1 and DIC-1 (Fig 3A). Together, these observations suggest that the Integrator plays a role in regulating the production or processing of piRNA precursors.

**Figure 3.**
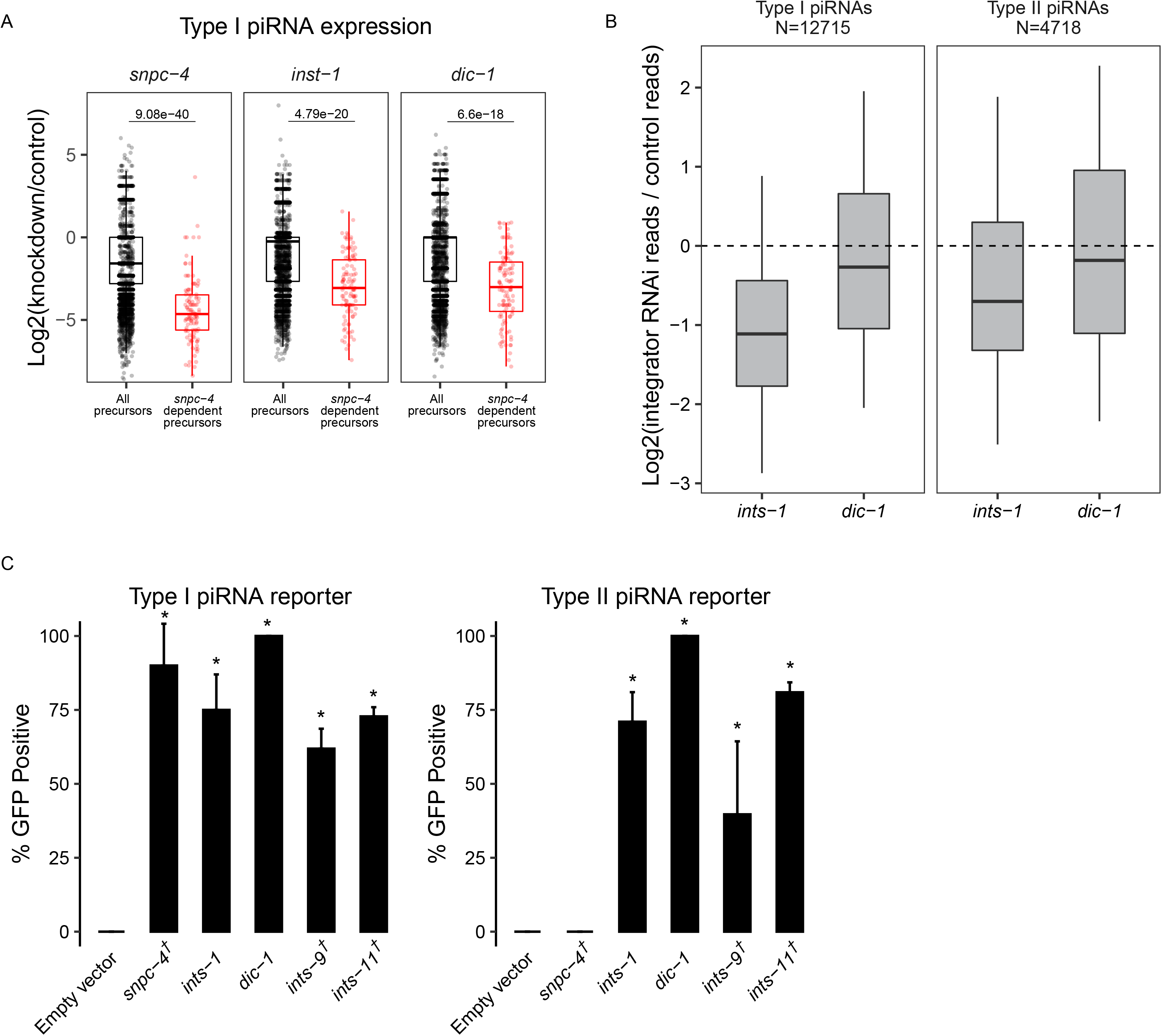
The Integrator complex promotes piRNA biogenesis. A) piRNA precursor fold changes in indicated knockdowns relative to empty vector control knockdown. All piRNA precursors are compared to precursors originating from loci which most depend on *snpc-4* for the accumulation of their mature sequences. B) piRNA fold changes in indicated knockdowns relative to empty vector control knockdown. Type I and type II piRNAs are compared separately. C) GFP expression in the type I and type II piRNA reporters in indicated RNAi knockdowns. † indicates L1 to adult same generation RNAi treatment. Asterisks indicate significant reporter activation, see Methods for details.

At snRNA loci in *C.elegans* and *Drosophila*, loss of the Integrator complex results in transcriptional readthrough and consequently elongated snRNA transcripts (11,27). Therefore, we predicted that Integrator depletion would result in readthrough of piRNA precursors at piRNA loci. Since piRNA precursor molecules are rare compared to mature piRNAs and other abundant small RNA species, we performed CapSeq to enrich for capped piRNA precursor molecules (28). In empty vector treated control animals, we found the size of piRNA precursor lengths peak at 25, 41, and 61 nucleotides. Following *ints-1* depletion, animals showed less defined peaks and elongated piRNA precursors (Fig S2C), consistent with a role for the Integrator complex in transcriptional termination of piRNA precursors. These findings support a model where at type I piRNA loci, SNPC-4 recruits a subset of the Integrator complex to elongating RNA pol II, leading to the endonucleolytic cleavage and subsequent termination of nascent piRNA precursors (Fig S2D).

SNPC-4 interacts with an upstream sequence called the Ruby motif, which is only found at type I piRNA loci (2). Surprisingly, we found that both type I and type II piRNA accumulation were globally reduced following *ints-1* and *dic-1* knockdown, although type II accumulation was less affected than type I following depletion of either component (Fig 3B). Our finding contradicts the recent report that INTS-11 does not function at type II piRNA loci (9), so we sought to clarify this point. To investigate whether the Integrator is required for gene silencing by type II piRNAs, we built a sensitized piRNA reporter where the GFP-targeting piRNA is made from a type II piRNA locus (28). We found that *snpc-4* depletion leads to only type I reporter activation but not type II reporter activation. However, RNAi depletion of *ints-1*, *dic-1, ints-9*, or *ints-11* resulted in GFP expression in both type I and type II reporters (Fig 3C). This suggests that at type I piRNA loci, the Integrator functions with SNPC-4 to promote piRNA accumulation while at type II piRNA loci, the Integrator can function in a SNPC-4 independent manner to promote piRNA biogenesis.

### Select nuclear pore and nucleolus components maintain CSR-1 perinuclear accumulation and promote piRNA silencing

Out of over 20 nuclear pore components tested, only 4 nuclear pore components (*npp-1*, *npp-6*, *npp-7*, and *npp-9*) caused over 40% GFP expression in our reporter when depleted (Fig 4A), despite most treatments resulting in morphologically effected worms (Supplementary Table 1).

**Figure 4.**
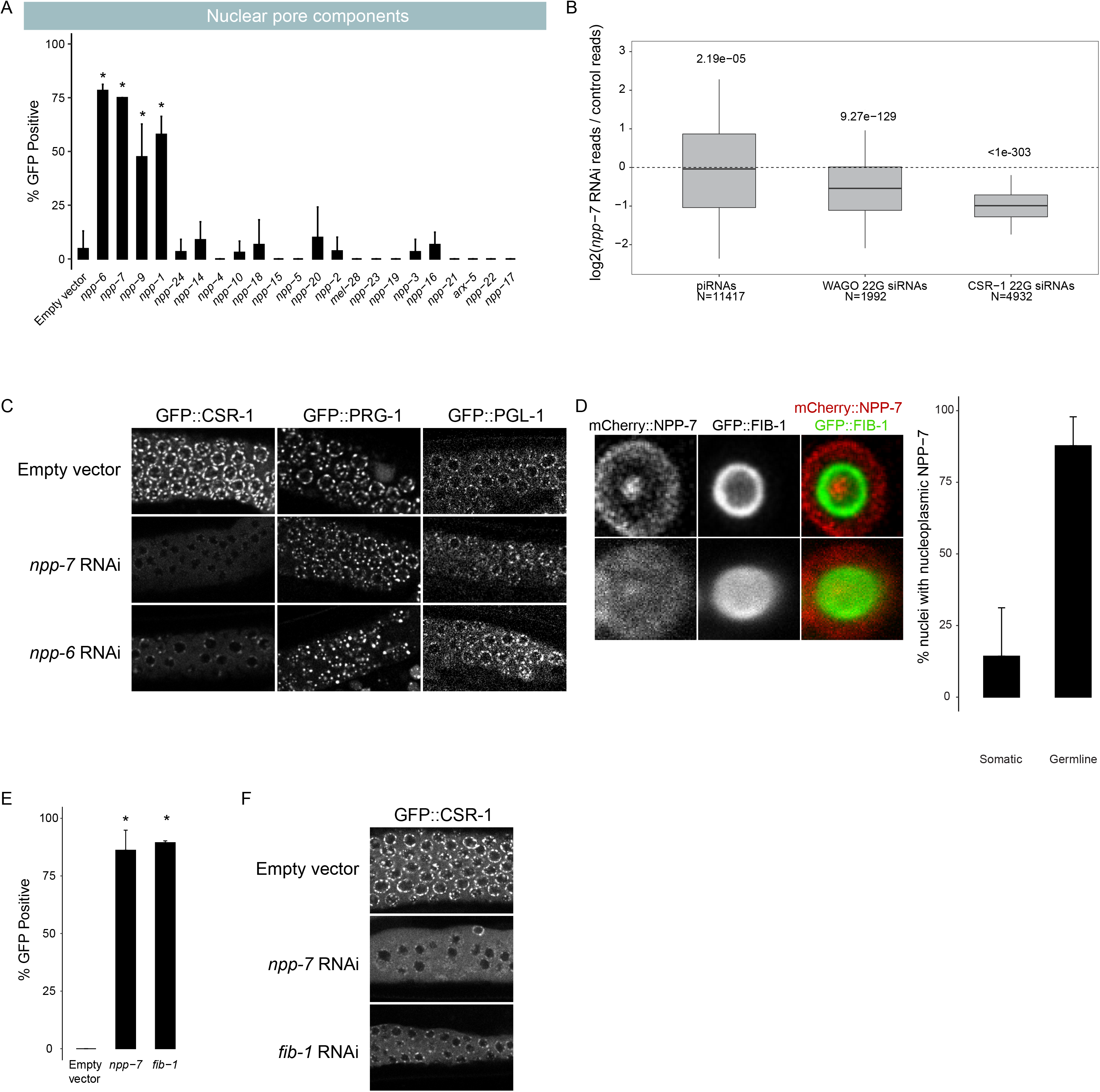
Nuclear pore component NPP-7 and the nucleolus promote piRNA dependent silencing. A) GFP expression in the piRNA reporter in indicated RNAi knockdowns. Asterisks indicate significant reporter activation, see Methods for details. B) Small RNA fold changes in *npp-7* RNAi knockdowns relative to empty vector control knockdown. C) Fluorescent micrographs of live adult worm gonads show the localization of GFP∷CSR-1, GFP∷PRG-1, and GFP∷PGL-1 in indicated RNAi knockdowns. D) Fluorescent micrograph of individual live adult worm germline nucleus shows the localization of mCherry∷NPP-7 and GFP∷FIB-1. Top panels display a single slice through the nucleus center, bottom panels display average projection of all slices though the full nucleus volume. Percentage of soma and germline nuclei with nucleoli that contain NPP-7 are quantified to the right. E) GFP expression in piRNA reporter in indicated RNAi knockdowns. Asterisks indicate significant reporter activation, see Methods for details. F) Fluorescent micrographs of live adult worm gonads show the localization of GFP∷CSR-1 in indicated RNAi knockdowns.

This suggests that only a subset of the factors that make up the nuclear pore are essential for piRNA silencing. As describe above, we found that loss of *npp-7* resulted in a reduction in 22G-RNAs but not in piRNAs, (Fig 2B), suggesting that *npp-7* is not critical for piRNA biogenesis but rather functions in the production of secondary WAGO 22G-RNAs. Furthermore, when we performed RNAi against *npp-1*, *npp-6*, or *npp-7* using the type II piRNA reporter established above, each treatment led to type II reporter activation (Fig S3A), consistent with nuclear pore components functioning downstream of piRNA biogenesis. Surprisingly, 22G-RNAs that map to CSR-1 targeted genes were significantly more reduced than 22G-RNAs that map to WAGO targeted genes following *npp-7* knockdown (Fig 4B), indicating that NPP-7 may play a critical role in regulating CSR-1 function.

CSR-1 is an Argonaute protein that prevents PRG-1 from targeting many endogenous germline genes (29). Consistent with a more pronounced loss of CSR-1 bound 22G RNAs following *npp-7* knockdown, we also observed mis-localization of CSR-1 protein in *npp-7* and *npp-6* RNAi treated animals (Fig 4C). These results suggest that the production and/or function of CSR-1 small RNAs are likely compromised in *npp-7* depleted animals. If the GFP expression that we observed in our piRNA reporter following *npp-7* knockdown was caused by a CSR-1 defect, then we reasoned that knockdown of *csr-1* itself by RNAi should also result in GFP expression. Indeed, we found that knockdown of *csr-1* and of the H3K36 methyl transferase *mes-4* both caused GFP expression in our reporter (Fig S3B). This suggests that disruption of CSR-1 or of chromatin modifiers associated with promoting gene expression can also impact piRNA mediated gene silencing. Therefore, our observations suggest that *npp-6* and *npp-7* may contribute to piRNA silencing by regulating the CSR-1 pathway.

We wondered how predictive dispersal of CSR-1 was on piRNA reporter activation for the remaining nuclear pore components. Therefore, we monitored the expression of CSR-1 and PRG-1 in live adult animals following knockdown of all previously screened nuclear pore components (Fig S3C). We found that knockdown of *npp-1* caused CSR-1 but not PRG-1 to become mis-localized as with *npp-7* and *npp-6*. These observations suggest that NPP-1, NPP-6, and NPP-7 specifically impact the localization of CSR-1 but not that of other P granule factors at the nuclear periphery, and these three factors also promote piRNA-dependent gene silencing.

On the other hand, *npp-9* knockdown led to piRNA reporter activation but did not cause dispersal of CSR-1 or of PRG-1. This result was surprising, as it has been previously shown that RNAi knockdown of *npp-9* does cause CSR-1 dispersal in *C.elegans* embryos (14). Conversely, knockdown of *npp-24* and *npp-2* cause CSR-1 dispersal but do not lead to piRNA reporter activation. Therefore, there is an imperfect correlation between nuclear pore component loss leading to piRNA reporter activation and CSR-1 dispersal. Nonetheless, our study reveals that select nuclear pore components, including NPP-1/6/7, promote CSR-1 perinuclear localization and are required for proper piRNA silencing.

The *Drosophila* homolog of NPP-7, Nup153, has been shown to function both at the nuclear membrane as a component of the nuclear pore and as a soluble component within the nucleoplasm to promote both gene expression and silencing (30,31). We therefore wondered whether NPP-7 in *C.elegans* may also be expressed intranuclearly to help CSR-1 define transcriptionally active regions of the genome. We used the CRISPR/Cas9 system to tag endogenous NPP-7 with mCherry and observed its expression in adult animals, and we found that in addition to its strong expression at the nuclear periphery, NPP-7 is also expressed as a single distinct focus in the nucleoplasm (Fig 4D). Surprisingly, we found that the single focus of intranuclear NPP-7 in germ cells resides in the nucleolus, and the constitutive nucleolar component FIB-1/fibrillarin surrounds the intranuclear NPP-7 focus. We found that nucleolar NPP-7 accumulates in nearly every germline nucleus but only rarely in somatic nuclei.

Additionally, nucleolar but not membrane associated accumulation of NPP-7 relies on FIB-1 (Fig S3D). We wondered whether reporter GFP expression following *npp-7* RNAi occurred due to loss of membrane-associated or nucleolar NPP-7. If nucleolar NPP-7 is required for piRNA silencing, then we expected that loss of FIB-1, which disrupts nucleolar NPP-7 accumulation, would cause piRNA reporter activation. Indeed, *fib-1* RNAi led to GFP expression in the piRNA reporter, consistent with involvement of the nucleolus in the piRNA pathway (Fig 4E). This result does not exclude the possibility that both nucleolar as well as membrane bound NPP-7 are mutually required for piRNA silencing. In vertebrae, the NPP-2 homologue Nup85 is required for nuclear pore assembly following cell division (32). We reasoned that depletion of NPP-2 might disrupt the membrane bound fraction of NPP-7 while leaving the nucleolar fraction unaffected, giving us an opportunity to observe the consequences of membrane bound but not nucleolar loss of NPP-7. Indeed, we found that *npp-2* knockdown affected the enrichment of NPP-7 in the nuclear membrane, resulting in a greater proportion of NPP-7 signal in the nucleolus (Fig S3D).

As shown above, *npp-2* knockdown also leads to CSR-1 dispersal from perinuclear granules. The presence of nucleolar NPP-7 may explain why *npp-2* knockdown does not compromise piRNA reporter silencing. Furthermore, *npp-1* and *npp-6* knockdown selectively disrupted NPP-7 nucleolar accumulation and also led to sensitized reporter GFP expression and CSR-1 dispersal. Nonetheless, the correlation of nucleolar NPP-7 localization and piRNA silencing was not perfect as *npp-5* knockdown disrupted nucleolar NPP-7 accumulation but did not lead to reporter GFP expression or CSR-1 dispersal. Nevertheless, this correlation suggests that nucleolar and not membrane bound NPP-7 is involved in the piRNA pathway.

We next sought to reconcile the two observations concerning NPP-7 presented here: (1) the involvement of NPP-7 in CSR-1 localization and function and (2) the role of nucleolar NPP-7 in piRNA pathway function. We hypothesized that these two observations may be directly connected if nucleolar NPP-7 is required for CSR-1 localization. To test this, we performed *fib-1* RNAi in a GFP∷CSR-1 tagged strain. Remarkably, *fib-1* knockdown disrupts perinuclear CSR-1 accumulation (Fig 4F), but not PRG-1 perinuclear accumulation similarly to *npp-7* knockdown (Fig S3D). Together, our study uncovers roles for select nuclear pore and nucleolus components in promoting CSR-1 localization and piRNA silencing.

### Importin components promote HRDE-1 silencing and localization

We found that depletion of 3 Importin family components, *ima-3*, *imb-2*, and *imb-5*, caused GFP expression in the sensitized reporter (Fig 2A). Activation during *ima-*3 and *imb-*5 knockdown also occurred using the type II piRNA reporter, supporting a role for the Importin family downstream of piRNA biogenesis (Fig S4A). Since knockdown of HRDE-1 cofactors NRDE-1 and NRDE-2 led to activation of our piRNA reporter, we hypothesized that HRDE-1 nuclear import might be compromised following depletion of these factors, which would in turn impact piRNA mediated silencing. To test this, we treated a GFP∷HRDE-1 tagged animal with *ima-3* and observed HRDE-1 localization in dissected adult gonads using confocal microscopy. Indeed, in the *ima-3* depleted germline, HRDE-1 failed to localize to nuclei. On the contrary, in the *ima-1* depleted germline, which continues to silence the piRNA reporter, HRDE-1 localization was unaffected (Fig 5A). These results are consistent with a model in which IMA-3 promotes piRNA-mediated gene silencing by translocating 22G-RNA-bound HRDE-1 from the cytoplasm to the nucleus where it can interact with nascent RNAs to transcriptionally silence piRNA targets. Interestingly, the nuclear Argonaute protein downstream of the piRNA pathway in *Drosophila*, Piwi, also relies on IMA-3 for translocation into the nucleus following piRNA binding (22).

**Figure 5.**
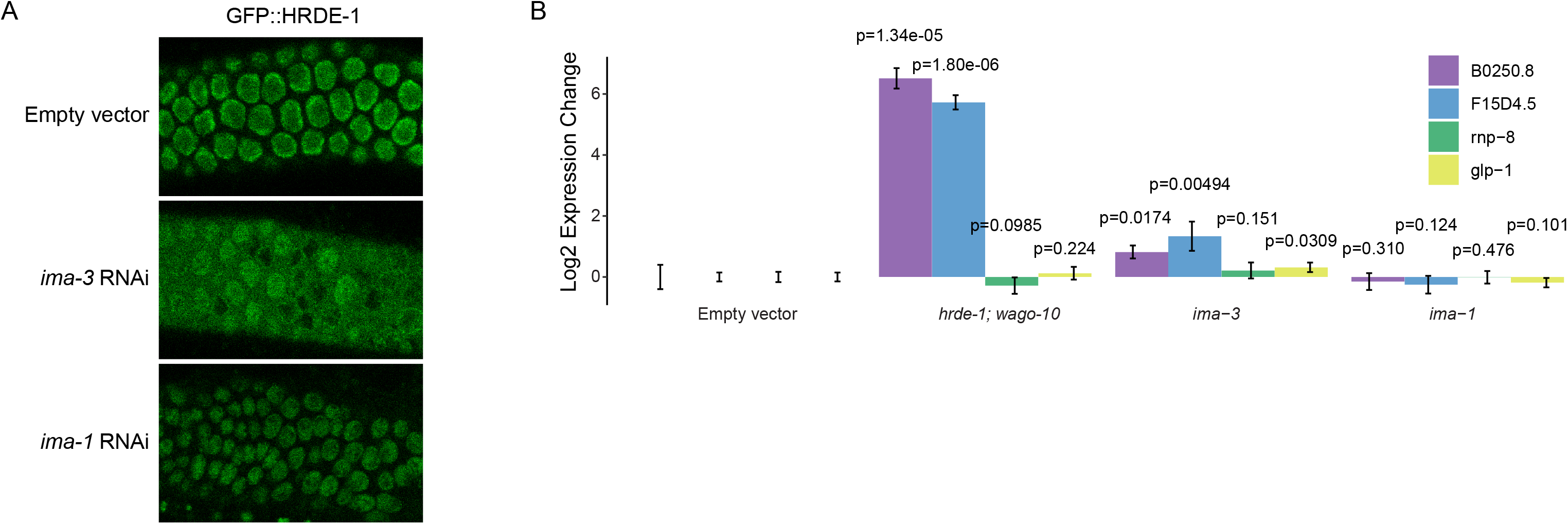
IMA-3 promotes HRDE-1 nuclear import. A) Fluorescent micrographs of live adult worm gonads show the localization of GFP∷HRDE-1 in indicated RNAi knockdowns. B) RT-qPCR results show expression change in indicated genotypes (*hrde-1; wago-10*) or RNAi knockdowns (Empty vector, *ima-3*, and *ima-1*). Statistical significance from two-tailed student’s t-test compared to control is displayed above treatments.

To further support this model, we asked whether endogenous genes that are silenced by HRDE-1 also rely on IMA-3 to maintain that silenced state. Using RT-qPCR, we found that *B0250.8* and *F15D4.5* are significantly upregulated both in *hrde-1* mutants and following *ima-3* RNAi treatment (Fig 5B). Therefore, we conclude that HRDE-1 translocates from the cytoplasm to the nucleus using the Importin α member IMA-3. Importin α members are known to function as adaptor molecules between protein cargo that contains a nuclear localization sequence (NLS) and Importin β members (33). In *C.elegans*, the somatic nuclear Argonaute protein NRDE-3 contains a NLS that, when mutated, results in failure of NRDE-3 to translocate into nuclei (23). Surprisingly, despite sharing significant sequence identity with HRDE-1, the NLS within NRDE-3 poorly aligns with HRDE-1 (Fig S4B). Therefore, how IMA-3 recognizes HRDE-1 for nuclear translocation requires further study.

## Discussion

We have utilized a sensitized reporter strain which detects defects in both piRNA biogenesis and effector factors to identify and characterize factors involved in piRNA silencing. Using our reporter, we have identified pre-mRNA splicing factors, Integrator complex subunits, protein import components, and nuclear pore components as essential for piRNA-mediated gene silencing. While we have currently used our reporter in a candidate screen, it is likely possible to use this strain in a forward screen, which would allow for even less biased detection of piRNA pathway components. Because the reporter robustly expresses germline GFP upon piRNA pathway disruption, activated worms containing a mutation in essential pathway components could possibly be sorted from silenced worms, for example using a COPAS BIOSORT system (34). Whether the expression of GFP can be distinguished from the significant auto-fluorescence contributed by intestinal granules adjacent to and overlapping with the adult gonad by a worm sorter remains to be seen and could represent a significant barrier to a high throughput approach.

We found that a subset of the Integrator complex contributes to both type I and type II piRNA biogenesis. Because both snRNA loci and type I piRNA loci require the SNAPc complex for biogenesis (25,26), we expected that the Integrator complex would be involved in piRNA biogenesis as described by recent reports (9,10). Our reporter assay and sequencing data support this model, but unexpectedly and inconsistent with the previously published findings, we also noticed that type II piRNA biogenesis and silencing were compromised following Integrator complex knockdown. Because SNPC-4 is recruited to Ruby motif-containing type I piRNA loci and not type II piRNA loci (2,35), our data suggest that the Integrator also functions at type II piRNA loci in a SNPC-4 independent manner. It has been shown that the Integrator can also function at protein coding genes to terminate transcription, promote RNAP II transcription proximal pausing, and promote enhancer RNA accumulation (36–40). While the canonical function of the Integrator at snRNA loci requires recognition of a conserved 3’ box sequence (41), protein-coding genes which rely on the Integrator complex to cleave nascent RNAs do not contain a recognizable 3’ box sequence (36). Intriguingly, *C.elegans* snRNA loci are devoid of a 3’ box sequence as well (42). Our data suggest that type II piRNA biogenesis represents yet another non-canonical role for the Integrator complex. Further investigation into the commonalities between snRNA loci, type I piRNA loci, type II piRNA loci, and protein coding genes may help to explain how the Integrator complex recognizes and cleaves nascent RNA molecules in *C.elegans*, and also may shed light on the Integrator’s non-canonical roles in other organisms.

We found that a nuclear pore component, NPP-7, localizes to both the nuclear envelope and the nucleolus, and that depletion of both NPP-7 and of nucleolar component FIB-1 both activate our piRNA reporter. This suggests a potential role for the nucleolus in the piRNA pathway. In *Drosophila* ovarian somatic cells, nuclear Argonaute Piwi has been shown to localize to nucleoli under heat shock, correlating with the expression of retrotransposons which have integrated into rDNA copies (43). Additionally, it has been shown in *C.elegans* that nuclear RNAi machinery targets rRNAs which are degraded by exosomes in the nucleolus (44). Both of these observations together with our own data suggest that the nucleolus may be involved in specialized small RNA-mediated silencing events. Additionally and possibly connected to nucleolar NPP-7, we found that NPP-7 promotes the localization of germ granule component CSR-1, but not germ granule components PRG-1 or PGL-1. We recently showed that CSR-1 uniquely retains its perinuclear localization when VASA helicases are mutated while PRG-1 and PGL-1 each show significant dispersal, suggesting that CSR-1 is recruited to the perinucleus in a VASA-independent manner (15). Our results suggest that the nuclear pore itself may represent a part of this VASA-independent pathway, and further that CSR-1 recruitment may require involvement of the nucleolus as *fib-1* knockdown similarly led to specifically CSR-1 dispersal.

We observed that knockdown of licensing Argonaute *csr-1* and of the H3K36 methyl transferase *mes-4* both caused GFP expression in our piRNA reporter (Fig S2B). It has been shown that H3K36me3 occupancy and CSR-1 22G RNA targeting are highly correlated (45). However, it has also been shown that CSR-1 does not directly induce H3K36 methylation (46). Therefore, we hypothesize that our piRNA reporter is activated when pathways that promote gene expression as well as those that control gene repression are disrupted. If correct, then this model would suggest that a balance between gene silencing and expression exists within the germline and disruption of pathways that control either gene silencing or gene expression can impinge on the alternative. Consistent with this notion, it has been shown that “resetting” germline RNAi pathways by re-introducing RNAi machinery to RNAi deficient *C.elegans* in the absence of piRNAs led to potent sterility due to the silencing of essential genes (47,48). This finding suggests that silencing machinery is essential for keeping pathways that control silencing and expression distinct from one another and balanced in the germline. Our finding, conversely, suggests that factors which promote gene expression are also essential for maintaining this balance.

Altogether, we used a piRNA reporter assay to identify and characterize factors that act at multiple levels of gene silencing, revealing a role for snRNA processing machinery, pre-mRNA splicing factors, the nuclear pore, the nucleolus, and protein import machinery in initiating and maintaining small RNA-mediated gene regulation (Fig 6).

**Figure 6.**
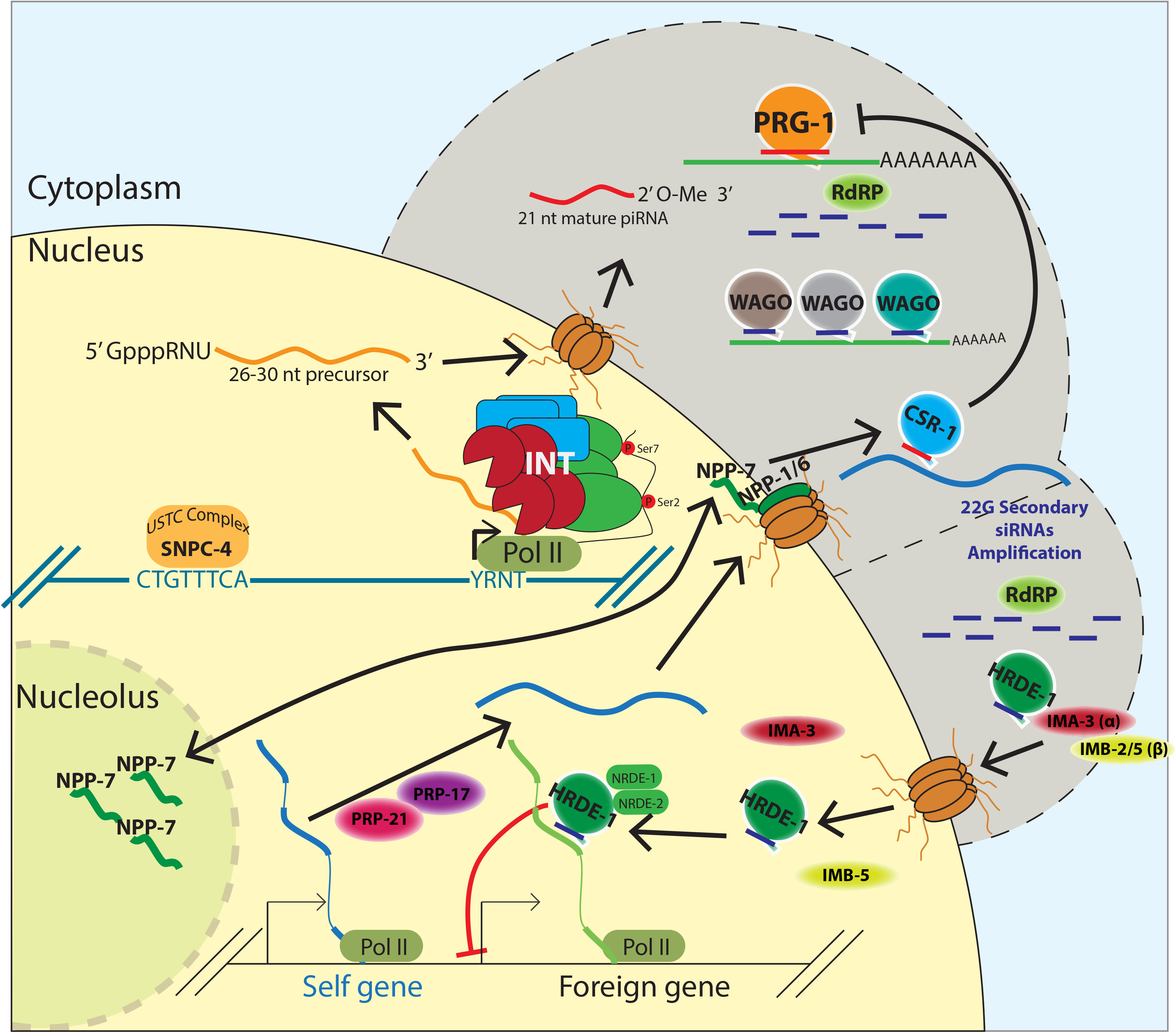
Multifaceted involvement of conserved and essential machinery work in concert with small RNA-mediated gene silencing in germ cells. A model depicting a role for factors identified in our screen. The Integrator complex promotes piRNA biogenesis, a subset of nuclear pore factors promotes CSR-1 localization and interface with the nucleolus, second step splicing factors ensure proper endogenous splicing occurs preferentially, and protein import machinery allow for the nuclear Argonaute HRDE-1 to translocate into the nucleus to transcriptionally silence target genes.

## Methods

### *C.elegans* strains

Animals were grown on standard nematode growth media (NGM) plates seeded with the *Escherichia coli* OP50 strain at 20°C. COP262 (*knuSi221 [fib-1p∷fib-1(genomic)∷eGFP∷fib-1 3’ UTR + unc-119(+)] II*.) and SS747 (*bnIs1[pie-1∷GFP∷pgl-1 + unc-119(+)]*) and YY538 (*hrde-1 (tm1200) III*.) were obtained from the Caenorhabditis Genetics Center (CGC). YY584 (*ggSi1[hrde-1p∷3xflag∷gfp∷hrde-1] II*.) was a gift from Scott Kennedy. HCL199 (*hrde-1 (tm1200) III; wago-10 (tm1186) V*.) was outcrossed from WM191. HCL105 (*gfp∷csr-1 IV*.) and HCL125 (*gfp∷prg-1 flag∷mCherry∷glh-1 I*.) are described in (49). Creation of HCL202 (*mCherry∷npp-7(uoc21) I*.) and HCL135 (*prg-1(uoc8) oma-1∷gfp I; cdk-1∷gfp II; 21ur-anti-gfp(type I) IV*.) are described below.

### RNA interference

RNA interference (RNAi) was performed by feeding animals with E. coli HT115 (DE3) strains expressing the appropriate double-stranded RNA (dsRNA). RNAi bacterial strains were obtained from the Ahringer C. elegans RNAi Collection (Source BioScience). Bacterial cultures were grown in Luria broth supplemented with 100 μg/ml ampicillin and 50 μg/ml tetracycline overnight at 37°C. Cultures were seeded on NGM plates containing 100 μg/ml ampicillin, 50 μg/ml tetracycline, and 1 mM IPTG and incubated at room temperature until dry. L4 hermaphrodites were picked onto the plates for feeding at 20°C and removed after 24 hours.

Adult progeny were screened. In instances where L4 treatment resulted in early next generation arrest, L1s were plated and screened as adults. Experiments where same generation screening was necessary are indicated in the relevant figure legend or data table. HT115 (DE3) expressing empty RNAi vector L4440 was used as the control. To assign significance to treatments in the piRNA reporter activation screen, a threshold of 20% activation was used as control treatment sometime resulted in spurious activation, but never resulted in over 20% naïve activation at 20°C for type I or type II piRNA reporters.

### CRISPR

#### Cas9/sgRNA constructs

We used the online tool sgRNA Scorer 2.0 (https://crispr.med.harvard.edu/) to design sgRNAs. The sgRNAs were cloned into pDD162 (50) by overlapping PCR using pDD162 as the PCR template and the appropriate primers.

Overlapping PCR products were inserted into pDD162 linearized with SpeI/BsrBI digestion by the seamless ligation cloning extract (SLiCE) method (51).

#### Donor constructs

To generate the flag∷mCherry∷NPP-7 donor construct, 500 bp upstream and 500 bp downstream of the *npp-7* TSS, and the flag∷mCherry coding sequences were amplified by PCR using N2 genomic DNA or plasmids containing flag∷mCherry as templates. PCR fragments were inserted into pUC19 linearized with HindIII/KpnI digestion by SLiCE. Silent mutations were introduced in guide RNA targeting sites by site-directed mutagenesis in donor constructs using Phusion High-Fidelity DNA Polymerases (Thermo Fisher Scientific).

### Fluorescence Imaging

GFP- and RFP/mCherry-tagged fluorescent proteins were visualized in living nematodes by mounting young adult animals on 2% agarose pads with M9 buffer (22 mM KH2PO4, 42 mM Na2HPO4, and 86 mM NaCl) with 10 mM levamisole. Fluorescent images used for localization studies were captured using a Zeiss LSM800 confocal microscope with a Plan-Apochromat 40X/1.4 Oil objective. Fluorescent images used for screening for GFP expression were captured using a Zeiss Axio Imager M2 compound microscope with a Plan-Apochromat 40X/1.4 Oil objective.

### Quantitative real-time PCR

1 μg of total RNA was reverse transcribed with SuperScript IV Reverse Transcriptase (Invitrogen) in 1x reaction buffer, 2U SUPERase-In RNase Inhibitor (Invitrogen), 0.5 mM dNTPs, and 2.5 μM random hexamers. Each real-time PCR reaction consisted of 3 μL of cDNA, 1 μM forward gene-specific primer and 1 μM reverse gene-specific primer. The amplification was performed using iTaq Universal SYBR Green Supermix (Bio-Rad) on the Bio-Rad CFX96 Touch Real-Time PCR Detection System. The experiments were repeated for a total of three technical replicates.

### Small RNA sequencing

#### Library preparation

Total RNA was extracted from whole animals of ~100,000 synchronized young adults. Small (<200nt) RNAs were enriched with mirVana miRNA Isolation Kit (Ambion). In brief, 80 μL (200-300 μg) of total RNA, 400 μl of mirVana lysis/binding buffer and 48 μL of mirVana homogenate buffer were mixed well and incubated at room temperature for 5 minutes.

Then 176 μL of 100% ethanol was added and samples were spun at 2500 x g for 4 minutes at room temperature to pellet large (>200nt) RNAs. The supernatant was transferred to a new tube and small (<200nt) RNAs were precipitated with pre-cooled isopropanol at −70°C. Small RNAs were pelleted at 20,000 x g at 4°C for 30 minutes, washed once with 70% pre-cooled ethanol, and dissolved with nuclease-free water. 10 μg of small RNAs were fractionated on a 15% PAGE/7M urea gel, and RNA from 17 nt to 40 nt was excised from the gel. RNA was extracted by soaking the gel in 2 gel volumes of NaCl TE buffer (0.3 M NaCl, 10 mM Tris-HCl, 1 mM EDTA pH 7.5) overnight. The supernatant was collected through a gel filtration column. RNA was precipitated with isopropanol, washed once with 70% ethanol, and resuspended with 15 μL nuclease-free water. RNA samples were treated with RppH to convert 22G-RNA 5’ triphosphates to monophosphates in 1x reaction buffer, 10U RppH (New England Biolabs), and 20U SUPERase-In RNase Inhibitor (Invitrogen) for 3 hours at 37°C, followed by 5 minutes at 65°C to inactivate RppH. RNA was then concentrated with the RNA Clean and Concentrator-5 Kit (Zymo Research). Small RNA libraries were prepared according to the manufacturer’s protocol of the NEBNext Multiplex Small RNA Sample Prep Set for Illumina-Library Preparation (New England Biolabs). NEBNext Multiplex Oligos for Illumina Index Primers were used for library preparation (New England Biolabs). Libraries were sequenced using an Illumina HiSeq4000 to obtain single-end 50 nt sequences at the University of Chicago Genomic Facility. *Analysis*. Fastq reads were trimmed using custom perl scripts. Trimmed reads were aligned to the *C.elegans* genome build WS230 using bowtie ver 1.2.1.1 (52) with options −v 0 --best -- strata. After alignment, reads that were between 17-40 nucleotides in length were overlapped with genomic features (rRNAs, tRNAs, snoRNAs, miRNAs, piRNAs, protein-coding genes, pseudogenes, transposons) using bedtools intersect (53). Sense and antisense reads mapping to individual miRNAs, piRNAs, protein-coding genes, pseudogenes, RNA/DNA transposons, simple repeats, and satellites were totaled and normalized to reads per million (RPM) by multiplying be 1e6 and dividing read counts by total mapped reads, minus reads mapping to structural RNAs (rRNAs, tRNAs, snoRNAs) because these sense reads likely represent degraded products. Reads mapping to multiple loci were penalized by dividing the read count by the number of loci they perfectly aligned to. Reads mapping to miRNAs and piRNAs were only considered if they matched to the sense annotation without any overlap. In other words, piRNA and miRNA reads that contained overhangs were not considered as mature piRNAs or miRNAs respectively. piRNA precursors were defined as sequences containing a full mature piRNA sequence plus a 2 nucleotide 5’ overhang corresponding to the genomic sequence 2 nucleotides upstream of that piRNA’s mature 5’ end. 22G-RNAs were defined as 21 to 23 nucleotide long reads with a 5’G that aligned antisense to protein-coding genes, pseudogenes, or transposons. RPM values were then used in all downstream analyses using custom R scripts using R version 4.0.0 (54), which rely on packages ggplot2 (55), reshape2 (56), ggpubr (57), dplyr (58).

### Cap-dependent RNA sequencing

#### Library preparation

Total RNA isolation was performed as described above for small RNA sequencing. Capped RNA molecules were enriched as described previously (28). Terminator exonuclease (Epicentre) was used to selectively degrade monophosphorylated RNAs. Quick CIP (New England Biolabs) was used to dephosphorylate non-capped triphosphorylated RNAs. RppH was used with Thermopol Buffer (New England Biolabs) to decap RNAs for ligation.

Libraries were prepared according to the manufacturer’s protocol of the NEBNext Multiplex Small RNA Sample Prep Set for Illumina-Library Preparation (New England Biolabs). NEBNext Multiplex Oligos for Illumina Index Primers were used for library preparation (New England Biolabs). Libraries were sequenced using an Illumina NovaSeq6000 to obtain single-end 100 nt sequences at the University of Chicago Genomic Facility.

#### Analysis

Adaptor trimming and alignment were performed as described above for small RNA sequencing. Reads that were between 17-100 nucleotides were retained. Precursor length histograms were constructed based on lengths of unique piRNA precursor sequences (molecules containing a full mature piRNA sequence plus the 2 nucleotides upstream of the mature piRNA 5’ end, as described above) from each library. Success of cap-dependent isolation was determined by comparing the enrichment for piRNA precursors versus mature piRNAs in the cap-dependent libraries compared to the previously constructed traditional small RNA libraries.

## Acknowledgements

This work is supported in part by NIH predoctoral training grant T32 GM07197 to J.B.; the NIH grant R01-GM132457 to H.-C.L.

