## Supplementary figures for "Sensitized piRNA reporter identifies multiple RNA processing factors involved in piRNA-mediated gene silencing"

**Title**

**This supplementary information includes:**

**Figure S1. piRNA reporter screening strategy.**

**Figure S2. Loss of piRNAs following *snpc-4*, *ints-1*, and *dic-1* knockdown.**

**Figure S3. NPP-1/6/7/9 and CSR-1 pathway disruption activates the sensitized piRNA reporter.**

**Figure S4. HRDE-1 does not share NRDE-3's nuclear localization signal.**

**Supplementary Table 1. Full list of candidates screened using piRNA reporter.**

**Figure S1. piRNA reporter screening strategy.**

A) A model depicts the cellular context of biogenesis and effector steps in piRNA silencing.

**Figure S2. Loss of piRNAs following *snpc-4*, *ints-1*, and *dic-1* knockdown.**

A) Scatterplots show the expression of mature type I piRNAs in the indicated RNAi knockdowns.

B) Scatterplots show the expression of type I piRNA precursors in the indicated RNAi knockdowns.

C) A histogram shows the distribution of type I and type II piRNA precursor sequence lengths in cap-enriched sequencing libraries from *ints-1* and empty vector RNAi treated worms.

D) A model shows the proposed action of the Integrator complex at piRNA loci.

**Figure S3. NPP-1/6/7/9 and CSR-1 pathway disruption activates the sensitized piRNA reporter.**

A) GFP expression in type II piRNA reporter in indicated RNAi knockdowns. Asterisks indicate significant reporter activation, see Methods for details.

B) GFP expression in type I piRNA reporter in indicated RNAi knockdowns. Asterisks indicate significant reporter activation, see Methods for details.

C) Percentage of worms with dispersed GFP::CSR-1 (top) or GFP::PRG-1 (bottom) granules in the indicated RNAi knockdowns. Asterisks indicate $p < 0.05$ in one-tailed Student’s t-test.

D) Percentage of germline nuclei with nucleoli that contain NPP-7 following the indicated RNAi knockdowns. Representative fluorescent micrographs show mCherry::NPP-7 localization from indicated treatments. Treatments that also activate the piRNA reported are marked with an asterisk.

**Figure S4. HRDE-1 does not share NRDE-3's nuclear localization signal.**

A) GFP expression in type II piRNA reporter in indicated RNAi knockdowns.

B) A selection from the Smith-Waterman local alignment of HRDE-1 and NRDE-3 amino acid sequences that includes the NRDE-3 nuclear localization signal.

**Supplementary Table 1. Full list of candidates screened using piRNA reporter.**

B) A model depicts the cellular context of biogenesis and effector steps in piRNA silencing.

### Figure S1

A

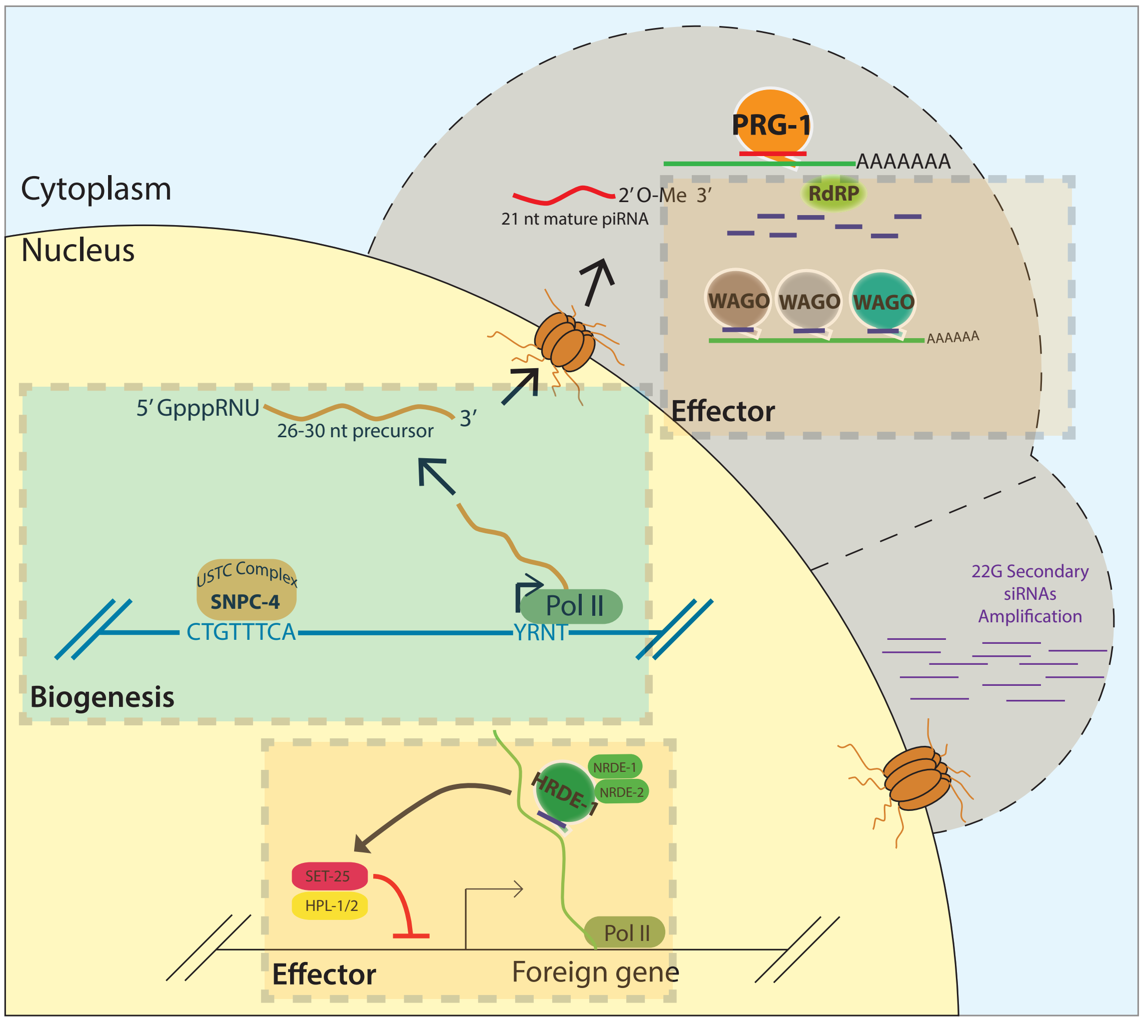

Cytoplasm
Nucleus
PRG-1
AAAAAAA
2′O-Me 3′
21 nt mature piRNA
RdRP
WAGO
WAGO
WAGO
AAAAAA
Effector
5′GpppRNU
26-30 nt precursor
3′
USTC Complex
SNPC-4
CTGTTTCA
Pol II
YRNT
Biogenesis
22G Secondary
siRNAs
Amplification
HRDE-1
NRDE-1
NRDE-2
SET-25
HPL-1/2
Pol II
Effector
Foreign gene

### Figure S2

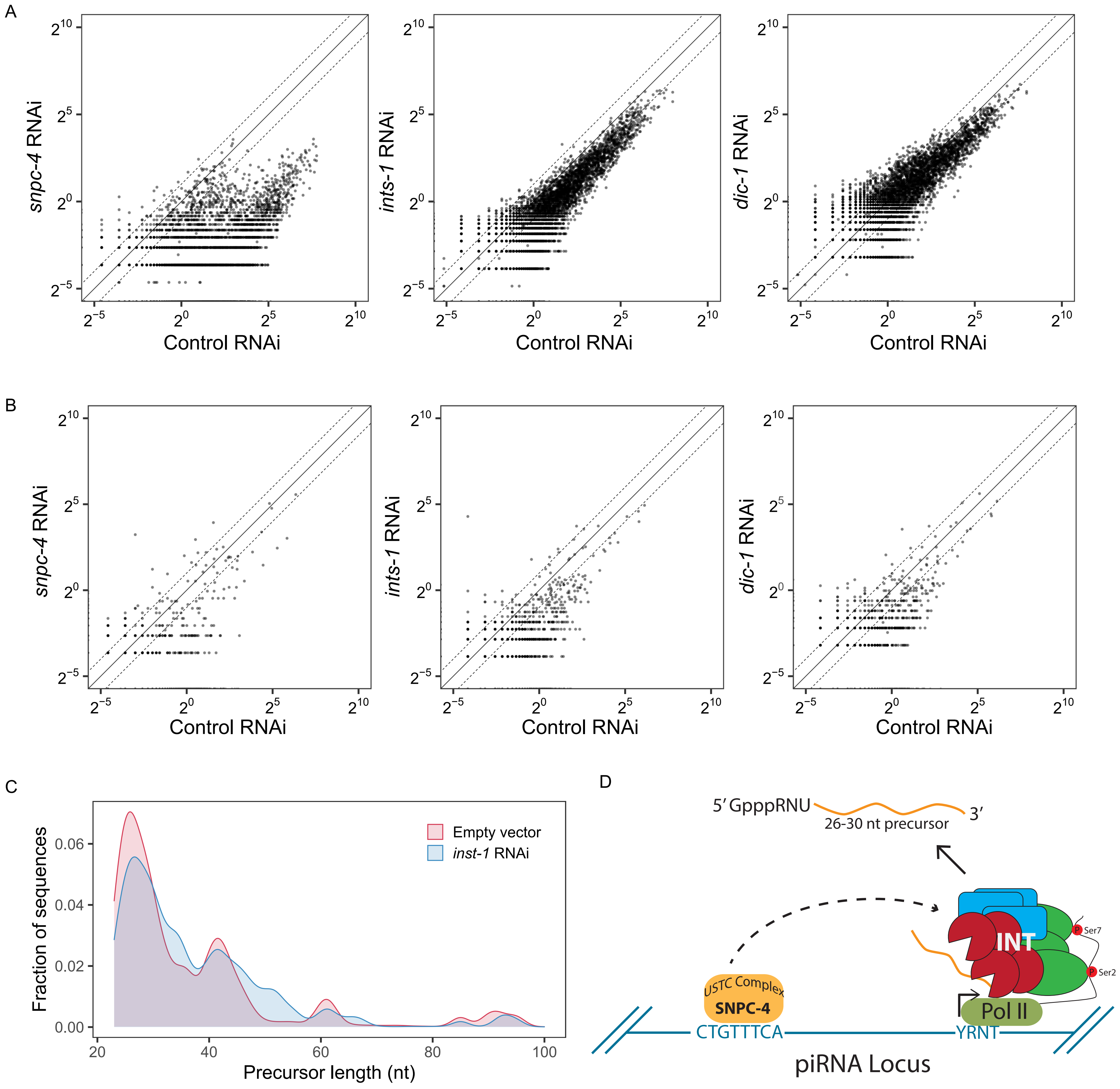

A
B
C
D
snpc-4 RNAi
ints-1 RNAi
dic-1 RNAi
Control RNAi
$2^{10}$
$2^{5}$
$2^{0}$
$2^{-5}$
Fraction of sequences
0.06
0.04
0.02
0.00
20
40
60
80
100
Precursor length (nt)
Empty vector
inst-1 RNAi
5′ GpppRNU
26-30 nt precursor
3′
INT
Ser7
Ser2
USTC Complex
SNPC-4
Pol II
CTGTTTCA
YRNT
piRNA Locus

### Figure S3

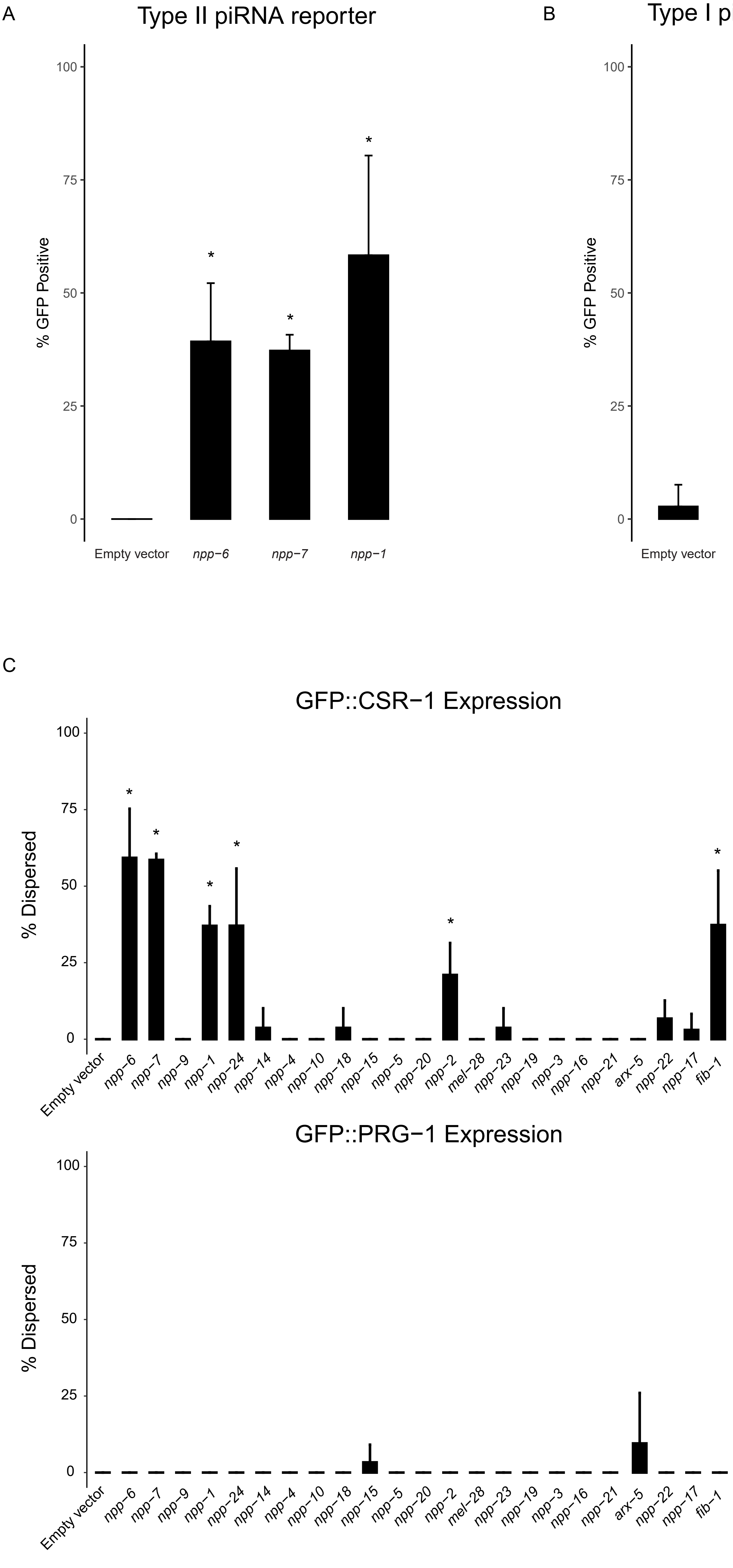

A
Type II piRNA reporter
% GFP Positive
100
75
50
25
0
Empty vector
npp−6
npp−7
npp−1
B
Type I piRNA reporter
% GFP Positive
Empty vector
mes−3
csr−1
C
GFP::CSR−1 Expression
% Dispersed
Empty vector
npp−6
npp−7
npp−9
npp−1
npp−24
npp−14
npp−4
npp−10
npp−18
npp−15
npp−5
npp−20
npp−2
mel−28
npp−23
npp−19
npp−3
npp−16
npp−21
arx−5
npp−22
npp−17
fib−1
GFP::PRG−1 Expression

### Figure S3, continued

D

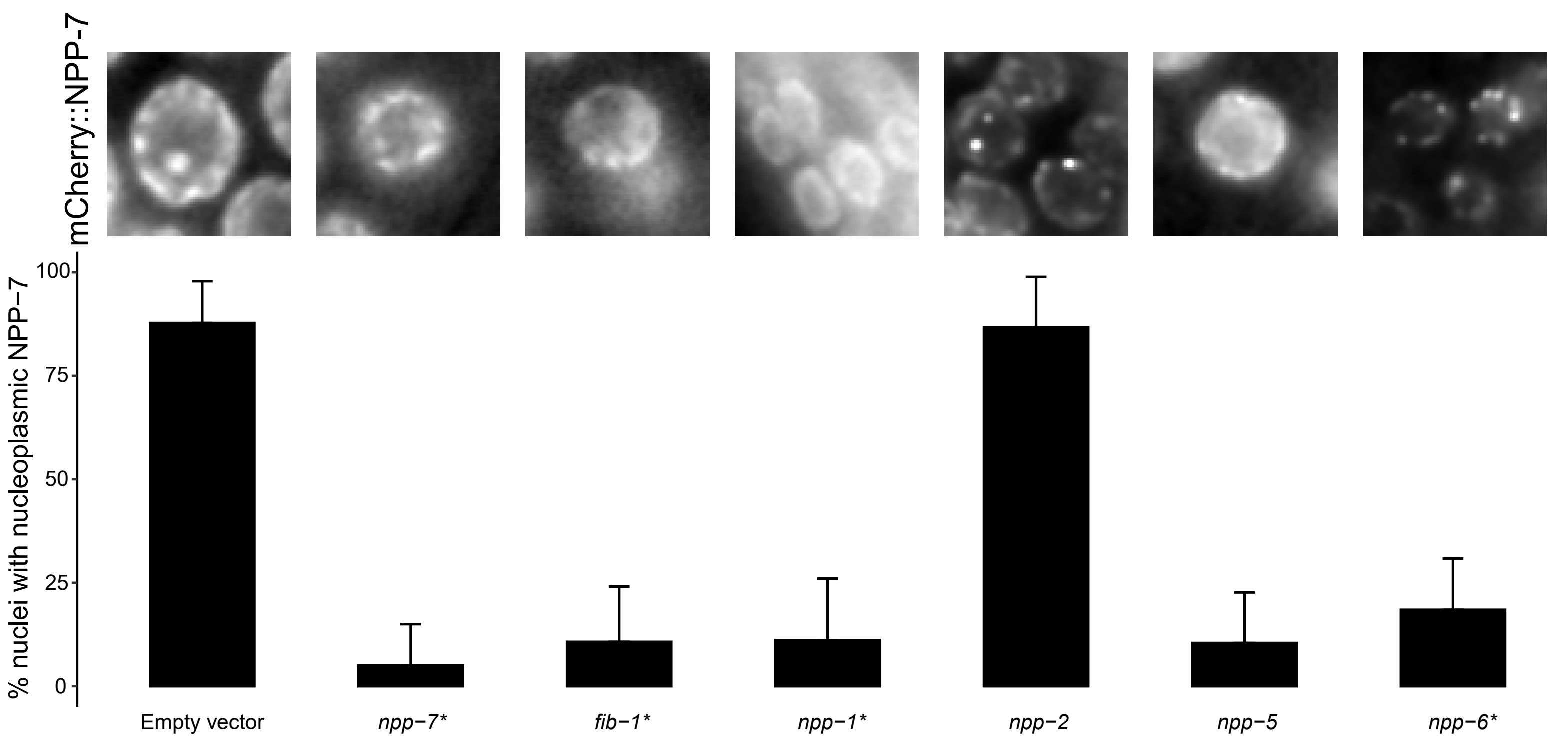


* = also activated piRNA reporter

### Figure S4

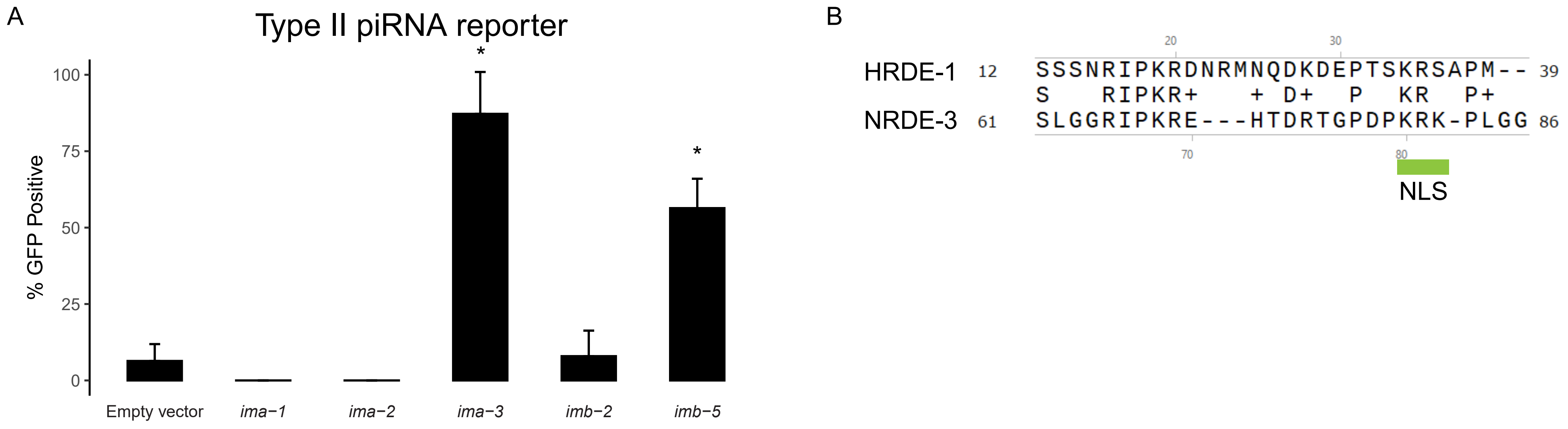
A
Type II piRNA reporter
% GFP Positive
100
75
50
25
0
*
*
Empty vector
ima−1
ima−2
ima−3
imb−2
imb−5
B
HRDE-1 12 SSSNRIPKRDNRMNQDKDEPTSKRSAPM-- 39
S RIPKR+ + D+ P KR P+
NRDE-3 61 SLGGRIPKRE---HTDRTGPDPKRK-PLGG 86
20
30
70
80
NLS
